# Parallel Computing to Speed up Whole-Genome Bayesian Regression Analyses Using Orthogonal Data Augmentation

**DOI:** 10.1101/148965

**Authors:** Cheng Hao, Garrick Dorian, Rohan Fernando

## Abstract

Bayesian multiple regression methods are widely used in whole-genome analyses to solve the problem that the number *p* of marker covariates is usually larger than the number *n* of observations. Inferences from most Bayesian methods are based on Markov chain Monte Carlo methods, where statistics are computed from a Markov chain constructed to have a stationary distribution equal to the posterior distribution of the unknown parameters. In practice, chains of about fifty thousand steps are typically used in whole-genome Bayesian regression analyses, which is computationally intensive. In this paper, we have shown how the sampling of marker effects can be made independent within each step of the chain. This is done by augmenting the marker covariate matrix by adding *p* new rows to it such that columns of the augmented marker covariate matrix are orthogonal. The phenotypes corresponding to the augmented rows of marker covariate matrix are considered missing. Ideally, the computations at each step of the MCMC chain, can be speeded up by the number *k* of computer processors up to the number *p* of markers. Addressing the heavy computational burden associated with Bayesian methods by parallel computing will lead to greater use of these methods.

## 2 Introduction

Genome-wide single nucleotide polymorphism (SNP) marker data have been adopted for whole genome analyses, including genomic prediction [9] and genome-wide association studies [13]. In whole-genome analyses, the number *p* of marker covariates is usually larger than the number *n* of observations. Bayesian multiple regression methods are widely used to address this problem, where the effects of all markers are estimated simultaneously combining the information from the phenotypic data and priors for the marker effects. Most widely-used Bayesian regression methods only differ in the prior used for the marker effects. For example, the prior for each marker effect in BayesA [9] follows a scaled t distribution, whereas several other Bayesian regression methods accommodate models where the prior for each marker effect follows a mixture distribution, such as BayesB [9], BayesC [7] and BayesR [2, 10].

In these Bayesian regression analyses, closed-form expressions for the posterior distribution of parameters of interest, e.g., marker effects, are usually not available. Thus inferences from most Bayesian methods are based on Markov chain Monte Carlo (MCMC) methods, where statistics are computed from a Markov chain constructed to have a stationary distribution equal to the posterior distribution of the unknown parameters. Suppose x is a stochastic vector of unknown parameters of interest. A Markov chain x_1_, x_2_, x_3_,…is a sequence of x, where the distribution of x*_t_* at step *t* conditional on all the previous steps only depends on the distribution of x*_t_*_−1_ at step *t* −1. It has been shown that statistics computed from such a Markov chain converge to those from the stationary distribution as the chain length increases [11]. In practice, chains of about fifty thousand steps are typically used in whole-genome Bayesian regression analyses [4]. Note that the vector x has length *p* or a multiple of it if auxiliary variables such as marker effect variances are introduced to the analysis as in BayesA or BayesB.

A widely used method to construct such a Markov chain is Gibbs sampling. In Gibbs sampling, at step *t*, each component of the vector x*_t_* is sampled from the conditional distribution of that component given all the other components sampled up to that point [12]. In a fast and efficient Gibbs sampler proposed for BayesB [1], for example, within each step, each variable in the vector x is sampled conditional on all the other variables. This includes, for each marker *i*, its effect, the effect variance and a Bernoulli variable indicating whether the effect is zero or non-zero, as well as the intercept and the residual variance. This is an example of a single-site Gibbs sampler where each variable is sampled at one time conditional on the current values of all other variables. In summary, whole-genome Bayesian multiple regression analyses require constructing Markov chains of length about fifty thousand. Within each step of the chain, Gibbs sampling requires sampling at least *p* unknowns. This makes Bayesian multiple regression analyses computationally intensive.

Parallel computing has been proposed to address this problem [14]. Parallel computing refers to the use of multiple processors to perform computations in parallel. It is often suggested that a large number of shorter chains can be constructed in parallel and combine the statistics computed from these chains. However, the Ergodic theorem of Markov chain theory states that statistics computed from an increasingly long chain, rather than an increasing number of short chains, converge to those from the stationary distribution [11]. Thus, combining several chains will reduce the Monte Carlo variance of the computed quantities, but this may not yield statistics from the stationary distribution. The problem with this approach is that a Markov chain is a sequential process, and thus it can not broken into several independent processes. However, a valid approach is to use Independent Metropolis-Hastings (IMH) sampling [12], where a large number of candidate samples x*_t_* are obtained independently using parallel computing. Then these candidate samples are accepted or rejected sequentially using the Metropolis-Hastings algorithm to construct a single long chain [8].

Another approach is to parallelize the Gibbs sampling for each marker within each step of the chain. In single-site Gibbs sampler, however, sampling of each variable is from the full conditional distribution, which is conditional distribution of the variable given the current values of all other variables. Thus, parallel Gibbs sampling would not be feasible unless the full conditional distributions do not depend on the values of the variables being conditioned on, i.e., unless the full-conditionals are independent. In this paper, we will show how the full conditional distributions of the marker effects can be made independent within each step of the chain. This is done by augmenting the marker covariate matrix by adding *p* new rows to it such that columns of the augmented marker covariate matrix are orthogonal [6]. The phenotypes corresponding to the augmented rows of marker covariate matrix are considered missing [6].

The computations for obtaining samples of the marker effects involves vector additions and dot products of length *n*. Parallel computing can also be used to speed up these computations, where vectors are split up and additions or products are done in parallel on multiple processors [5]. This approach can be used within each parallel Gibbs sampling.

The objective of this paper is to show how the parallel Gibbs sampling approach using an augmented marker covariate matrix can be used in Bayesian multiple regression methods with the BayesC prior. Use of this approach with other priors, such as those in BayesA, BayesB or Bayesian Lasso, should be straightforward.

## 3 Methods

### 3.1 Model

In Bayesian regression, phenotypes of are often modeled as

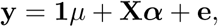

where y is the vector of *n* phenotypes, *μ* is the overall mean, **X** is the *n* × *p* marker covariate matrix (coded as 0, 1, 2), *α* is a vector of *p* random marker effects and e is a vector of *n* random residuals. A flat prior is used for *μ*. The prior for the residual e is 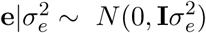 with 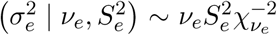. The columns of **X** are usually centered. In BayesC, the prior for the marker effect is a mixture of a point mass at zero and a univariate normal distribution with null mean and a common locus variance 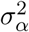 with 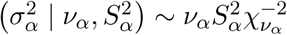 [7].

### 3.2 Parallel computing strategy using orthogonal data augmentation

#### 3.2.1 Gibbs sampling for marker effects in BayesC

In Gibbs sampling for BayesC, the full conditional distribution of *α_j_*, the marker effect for locus *j*, when *α_j_* is non-zero, can be written as

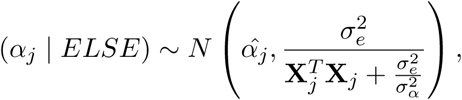

where *ELSE* stands for all the other unknowns and **y, X***_j_* is the jth column of **X,** and 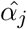 is the solution to
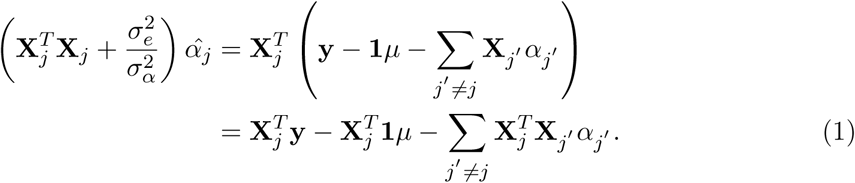

In the Gibbs sampling, the sample for each marker, *α_j_*, can not be obtained simultaneously in parallel, because samples for other marker effects, 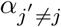, appear in the term 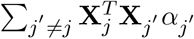, on the right-hand-side of (1), i.e., the full conditional distributions of the marker effects are not independent. One solution is to orthogonalize columns of the marker covariate matrix **X** such that the term 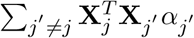 in (1) becomes zero. The data augmentation approach that is described below was proposed by Ghosh et al. [6] to obtain a design matrix with orthogonal columns.

#### 3.2.2 Orthogonal Data Augmentation (ODA)

Let **W***_o_* = [1 **X**] be the design matrix for the BayesC analysis. Following Ghosh et al. [6], we show here how to augment **W***_o_* as 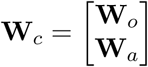 such that

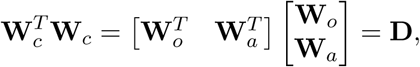

where **W***_a_* is a square matrix of dimension *p* and **D** is a diagonal matrix. Thus,
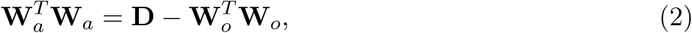

and **W***_a_* can be obtained using Cholesky decomposition (or Eigen decomposition) from (2). The choice of **D** is **I***d*, where *d* is set to be the largest eigenvalue of 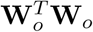 [6]. In practice, a small value, e.g., 0.001, was added to *d* to avoid computationally unstable solutions [6].

#### 3.2.3 BayesC model with ODA (BayesC-ODA)

Employing ODA, the Bayesian regression model can be written as
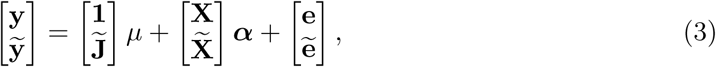

where 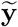 denotes a vector of unobserved phenotypes that are introduced into the model, 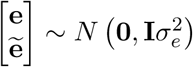 and 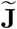, 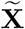 are obtained using (2) with 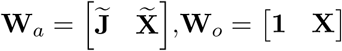,**W***_o_* = [1 **X**].

In BayesC-ODA, the full conditional distribution of ***α*** under model (3), which was derived in the Appendix, can be written as
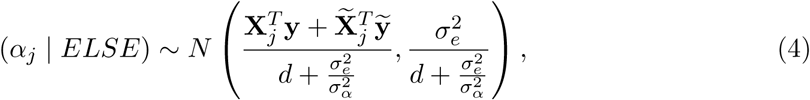

where the mean and variance parameters are free of the values of the other marker effects 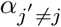. Thus the full conditional distribution of the marker effects are independent, and thus, samples for each marker can be obtained simultaneously in parallel. At each step of the MCMC chain, the “missing” phenotypes 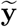 are sampled from
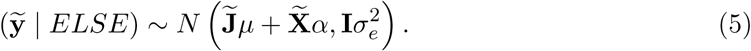

The derivation of the full conditional distributions of other parameters of interest are shown in the Appendix.

#### 3.2.4 Simulated data

Simulated genotypic and phenotypic data were used to compare BayesC and BayesC-ODA. The simulated genome consisted of 10 chromosomes each 5 cM long and containing 50 evenly spaced loci. Allele states were sampled from a Bernoulli distribution with frequency 0.5. A random sample of 25 loci were selected as QTL, and their effects were sampled from a univariate normal distribution with mean zero and variance one. Starting from a base population of 100 males and 100 females, random mating was simulated for 100 generations to generate linkage disequilibrium. In generation 101, the population size was increased to 3000 males and 3000 females, and random mating was continued for four more generations. The QTL effects were scaled such that the genetic variance for a randomly sampled individual from generation 105 was 1.0. Phenotypes were simulated by adding independent residuals that were sampled from a normal distribution with null mean and variance one to the genetic values. To investigate the performance of BayesC-ODA with *n* < *p* or *n* > *p*, 100 or 5000 individuals were used for training. A population of 1000 individuals was used for testing. In the testing population, estimated breeding values were calculated using BayesC and BayesC-ODA. Correlation between estimated breeding values or estimated marker effects from BayesC-ODA and BayesC was investigated for a chain of length 5,000,000 to study: 1) whether BayesC-ODA provided identical estimated marker effects and breeding values as BayesC; 2) the convergence of BayesC-ODA.

The true genetic variance and residual variance were used to calculate the scale parameters of the inverse-chisquare priors of the residual variance and marker effect variance [3].

## 4 Results

The correlation between estimated breeding values for the testing population from BayesC and BayesC-ODA by chain length was investigated. In the scenario where *n* < *p*, this correlation was larger than 0.99 when the chain was longer than 9,000 and became larger than 0.999 as the chain grew longer than 75,000. In the scenario where *n* > *p*, this correlation was larger than 0.99 when the chain was longer than 1,000 and became 0.999 as the chain grew longer than 18,000.

The correlation between posterior mean of marker effects from BayesC and BayesC-ODA as the chain length increases was investigated. In the scenario where *n* < *p*, this correlation was larger than 0.99 when the chain was longer than 37,000 and became larger than 0.999 as the chain grew longer than 439,000. In the scenario where *n* > *p*, this correlation was larger than 0.99 when the chain was longer than 649,000 and became about 0.999 as the chain length reached 5,000,000.

## 5 Discussion

Whole-genome Bayesian multiple regression methods are usually computationally intensive, where a MCMC chain of about fifty thousand steps is typically used for inference. In this paper, a strategy to parallelize Gibbs sampling for each marker within each step of the MCMC chain was proposed. This parallelization is accomplished by using an orthogonal data augmentation strategy, where the marker covariate matrix is augmented by adding *p* new rows such that its columns are orthogonal [6]. Then, the full conditional distributions of marker effects become independent within each step of the chain, and thus, samples of marker effects within each step can be drawn in parallel. In this paper, the full conditional distributions that are needed for BayesC with orthogonal data augmentation (BayesC-ODA) were derived and the convergence of BayesC-ODA was studied. In analyses of the simulated data, BayesC-ODA provided virtually identical predictions of breeding values as BayesC when the chain length was about 20,000 to 80,000, which is similar to the commonly used chain length of 50,000. Some ideas for parallel implementation of BayesC-ODA are briefly discussed below with more details in the appendix. The investigation of these ideas and parallel implementation of Bayesian multiple regression with ODA will be undertaken in a separate study.

In Bayesian multiple regression methods such as BayesC, the most time consuming task is sampling the marker effects from their full conditional distributions. In BayesC-ODA, however, the marker effects within each step can be sampled in parallel, using (4). Ideally, the computations at each step of the MCMC chain, can be speeded up by the number *k* of processors up to the number *p* of markers. However, two extra computations are required in BayesC-ODA. The first is sampling of the vector 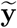 of unobserved phenotypes, which is required in each MCMC step. Each element of 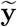 is sampled from an independent univariate normal distribution with the variance equal to the current value of 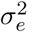. The means of these normal distributions can be computed in parallel as described in the appendix. Once the means are computed, each element in 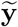 can be sampled in parallel. The second is the computation of the augmented matrix **W***_a_* as in (2), which is required only once at the beginning of the MCMC chain. In (2), there are two computationally intensive tasks: 1) computation of **X***^T^***X**, where **X** is a *n* × *p* matrix; and 2) Cholesky decomposition of a positive definite matrix of size *p*. Parallel computing approaches for the first of these two tasks is given in the appendix. The computing time for the Cholesky decomposition in the second task is relatively short, taking only a few minutes for *p* = 50, 000 on a workstation, using one graphics processing unit (GPU).

It is worth noting that two approaches are available to compute the right-hand-side of (1). In the first approach, equation (29) in [5] is used, where number of operations is of order *n*. In the second approach, equation (33) in [5] is used, where the number of operations is of order *p*. In BayesC-ODA, the first approach is used. As can be seen from (1), the right-hand-side for *α_j_* is 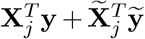, where 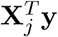 is constant, and only 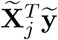 needs to be computed at each step of the MCMC chain, where the number of operations for this is always of order *p* regardless of the size of *n*. However, when the first approach is used for multiple-trait BayesC analyses, the size of the dataset that can be analyzed is limited by the requirement to store the entire marker covariate matrix of size *n* × *p* in memory so that **y** −1*μ* − ∑ **X***_j_α_j_* can be updated with the current value of *α_j_*. So, as *n* grows, this approach will become infeasible. On the other hand, in BayesC-ODA, only 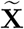 of constant size *p* × *p* needs to be stored in memory regardless of the size of *n*, which is required in (4) and (5). Thus, even when *n* grows, multiple-trait analyses will only require storing a *p* × *p* matrix regardless of the number of traits and *n*.

We have shown here that the predictions of breeding values from BayesC-ODA converge to those from BayesC but may require a chain of 80,000 steps as opposed to one of 50,000 for BayesC. However, Gibbs sampling of marker effects within each step can be done in parallel for BayesC-ODA, and this is expected to result in a considerable speedup for BayesC-ODA. Further, as discussed above, multiple-trait analyses with BayesC-ODA only require storing the *p* augmented rows of the covariate matrix regardless of the number of traits and observations. Thus, when *n* is large, BayesC-ODA may provide an efficient approach for multiple-trait Bayesian regression analyses.

## Author’s contributions

HC, RF contributed to the development of the statistical methods. HC wrote the program code and conducted the analyses. The manuscript was prepared by HC and RF. All authors read and approved the final manuscript.

## 6 Appendix

In many modern programming languages, such as R, Python and Julia, libraries are available to take advantage of multiple processors and GPUs for parallel computing of many matrix or vector operations. The descriptions given below are only to illustrate the main principle underlying parallel computing of splitting up calculations across processors. Actual implementations may be different and will depend on the programming language, the library and the hardware used.

### 6.1 Parallel Computing of Ab

To sample the unobserved phenotypic values using (5), a matrix by vector product 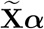 is needed. Here we describe how parallel computing can be used to compute the product of a matrix **A** by a vector **b**.

1. Split **A** of size *n* × *p* by columns into smaller matrices **A**_(1)_, **A**_(2)_, **A**_(3)_,…of size *n* × *p_i_*, and split *α* into smaller vectors **b**_(1)_, **b**_(2)_, **b**_(3)_,…of length *p_i_* with ∑ *p_i_* = *p*.
2. Compute **Ab** as **A**_(1)_**b**_(1)_ + **A***_j_*_(2)_**b**_(2)_ + **A***_j_*_(3)_**b**_(3)_ +…, where **A**_(*i*)_**b**_(*i*)_ for *i* = 1, 2,…are computed on different processors and then summed to obtain **Ab.**

The same strategy can also be used to calculate **X***^T^***y** by splitting **X** by rows.

### 6.2 Parallel Computing of A*^T^*A

In (2), computation of **X***^T^***X** is needed. Here we describe how parallel computing can be used to compute **A***^T^***A,** where **A** is a *n* × *p* matrix.

1. Split **X** of size *n* × *p* by rows into smaller matrices **A**_(1),_ **A**_(2)_, **A**_(3)_,…of size *n_i_* × *p* with *∑ n_i_* = *n*.
2. Compute 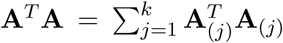, where 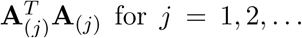 for *j* = 1, 2,…are computed on different processors and then summed to obtain **A***^T^***A.**

In addition to reducing the computing time, this approach can also address the limitation that **A** may be too large to be stored on a single computing node by distributing the **A**_(*i*)_ across several nodes.

### 6.3 Single-site Gibbs sampler for BayesC-ODA

#### 6.3.1 full conditional distribution of the marker effect

Detailed derivation of the full conditional distributions of the marker effect for locus *j* in BayesC is in Fernando and Garrick [3]. As shown in [3], the full conditional distribution of *α_j_* in BayesC, when *α_j_* is non-zero, is
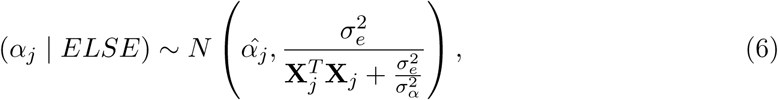

where *ELSE* stands for all the other unknowns and **y, X***_j_* is the *j*th column of **X,** and 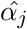 is the solution to
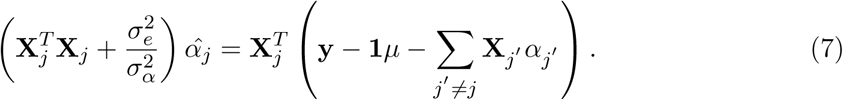

The full conditional distribution of *α_j_* in BayesC-ODA,which is shown below, can be obtained from (6) and (7) by replacing **y** with 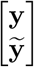, **1** with 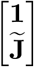 and **X** with 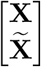. Note that columns of the augmented covariate matrix 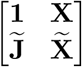 are orthogonal. Thus, (7) for BayesC-ODA can be simplified as
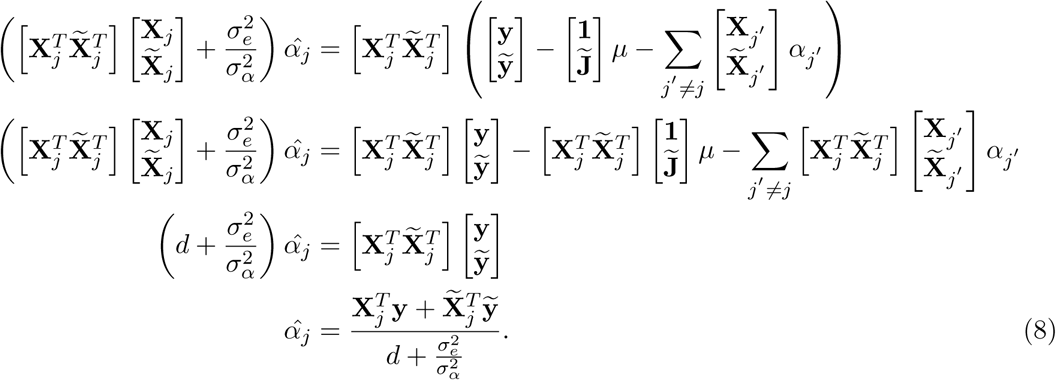

Thus, the full conditional distribution of *α_j_* can be written as

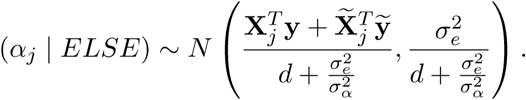

Detailed derivation of the full conditional distribution of the indicator variable *δ_j_* indicating if *α_j_* had a normal distribution (*δ_j_* = 1) or if it is null (*δ_j_* = 0) in BayesC is also in Fernando and Garrick [3]. The full conditional distribution of *δ_j_* in BayesC is
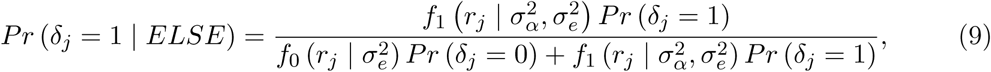

where 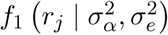 is a univariate normal with

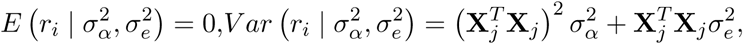

and 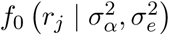 is a univariate normal with

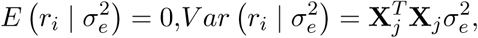

and

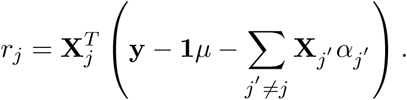

The full conditional distribution of *δ_j_* in BayesC-ODA, which is shown below, can be obtained from (9) by replacing **y** with 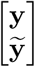, **1** with 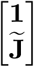 and **X** with 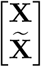. Thus, (9) for BayesC-ODA can be simplified as

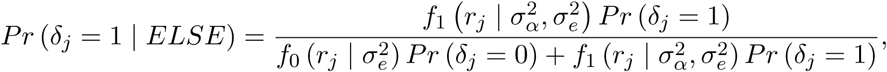

where 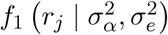 is a univariate normal with

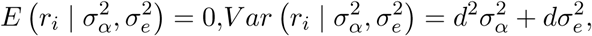

and 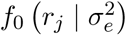 is a univariate normal with

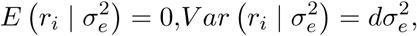

and

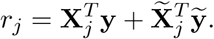

#### 6.3.2 full conditional distributions of the unobserved phenotypes

The full conditional distribution of 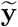 can be written as

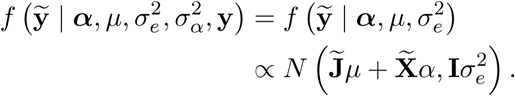

#### 6.3.3 full conditional distributions of other unknowns

The derivation of the full conditional distributions of other parameters such as *μ*, 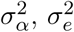 are straightforward. Thus they are presented as below without derivations.

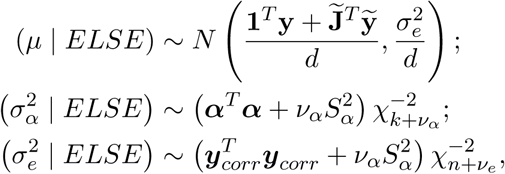

where 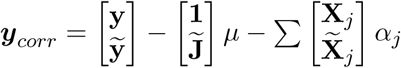 and *k* is the number of markers in the model.

